# Exploring cardiac phenotypes of Drosophila orthologues of human genes associated to Brugada Syndrome (BrS)

**DOI:** 10.1101/2025.07.08.663711

**Authors:** Sallouha Krifa, Julien Barc, Nathalie Gaborit, Flavien Charpentier, Laurent Perrin, Nathalie Arquier

## Abstract

Brugada syndrome (BrS) is a rare cardiac arrhythmic disorder with high risk of sudden cardiac death. Recent advances have identified more than 20 risk loci with complex inheritance suggesting a polygenic model for BrS inheritance. These loci are in non-coding regions located in the vicinity of cardiac-expressed genes. This complex genetic architecture and the limited understanding of BrS genetic and molecular mechanisms hinder the development of efficient prevention strategies in the context of this syndrome and are unfavorable to the implementation of therapeutic interventions. In this context, understanding the functional impact of the identified putative risk alleles is a prerequisite. Here, we used the fly model to systematically test whether orthologues of genes located near risk alleles for BrS participate to cardiac function. The fly is the simplest model with a heart muscle and is a powerful genetic model suitable for efficient screening of candidate genes, providing a whole organism-based assessment of cardiac development, structure and function. Using high-speed heart imaging platform on intact flies, we invalided the cardiac expression of the fly orthologues of human genes associated to BrS and characterized whether they are cell autonomously implicated in heart functioning. Our results provide an overview of cardiac phenotypes associated with genes potentially involved in BrS, enabling their prioritization for further investigations in mammalian models.

## INTRODUCTION

Sudden Cardiac Death (SCD) is a prominent cause of death in general population. Among the genetic causes of SCD, around 4-12% is due to Brugada syndrome (BrS)^1^. BrS is an inherited cardiac pathology, which affects mostly young adults with no clinical evidence of cardiac structural defects, that makes it difficult to detect and diagnose. An abnormal electrocardiogram (ECG) profile, with an ST-segment elevation and T-wave inversion in the right precordial leads, is the sole characteristic associated to BrS to date. Implantation of a defibrillator (ICD) is the only therapy proposed to avoid deleterious consequences of severe arrhythmias and SCD in patients^2^. However, this invasive approach has numerous side-effects, reinforcing a real need to find new ways to more deeply understand the pathology of BrS in order to find adapted prophylaxis or therapies. Indeed, the large gap in our knowledge concerning the determinants of BrS at the genetic and molecular levels hinders breakthroughs in the development of risk stratification and preventive strategies.

Long regarded as a monogenic disease with dominant autosomal transmission, recent studies showed that BrS onset is more complex than just resulting from a single causal gene^3^. Indeed, while 2-25% of BrS patients have rare coding variants in the SCN5A gene encoding the sodium channel Na _v_ 1.5 - whose dysfunctions has been associated to severe arrhythmia and SCD in BrS^4^ - these mutations do not explain the prevalence of BrS in the whole population. Indeed, more than 70% of BrS cases do not exhibit SCN5A variants nor other rare variants. These observations tend to suggest a polygenic origin of the disease, and calls for a deeper genetic analysis.

Thanks to the recruitment of a large cohort of BrS patients worldwide, two consecutive genome-wide association studies (GWAS) have been performed, leading to the identification of 21 common risk alleles, with no association to SCN5A rare variants^5,6^. The genetic variations associated to these alleles are located in non-coding regions, probably involved in gene expression regulation^5^. In addition, ten of them are located in close proximity of eight transcription factors (TF) known to be involved in cardiac development and physiology, but with one exception^7^, not to BrS disease. Three-dimensional structure analysis of the genome using Hi-C experiments performed on human left ventricles identified several loci physically interacting with the risk alleles, allowing to establish a list of 56 candidate genes that could be functionally linked to the onset of Brugada syndrome^5^.

Here we used the fly model to evaluate the potential cardiac function of these 56 genes. Drosophila is the simplest animal model with a beating heart, which is a monolayer tubular organ constituted of 84 cardiomyocytes^8^. The fly is increasingly used to identify genes involved in cardiac function in physiological and pathophysiological contexts, such as congenital heart diseases, ageing or metabolic cardiomyopathies^9–14^. It has also been used to investigate the genetic architecture of natural variations of heart function and heart aging^9,10^ and to evaluate the function of orthologues of genes located at loci identified by GWAS for cardiac disorders or heart rate regulation in human populations^15,16^. Drosophila heart is therefore a powerful genetic model suitable for efficient screening of gene candidates and providing a whole organism-based assessment of cardiac development, structure and function. In addition, the fly model is ideal for elucidating conserved genetic and molecular pathways because of low genetic redundancy and relative simplicity of genetic networks, thus allowing for efficient identification of novel genes that can provide insights into the genetics and pathogenesis of cardiac disorders. Furthermore, recent advances have established the Drosophila heart as a relatively high-throughput platform compared with other *in vivo* models. Using our high-speed heart imaging platform on whole flies, we therefore invalided the cardiac expression of the fly orthologues of the human genes associated to BrS risk alleles and characterized whether they are cell autonomously implicated in heart functioning. We characterized resulting cardiac phenotypes in one week old flies of both sexes. In order to prioritize genes for further analysis in mammalian models, we identified different categories of genes depending on the effect of their knock down on cardiac function in flies.

## RESULTS AND DISCUSSION

Our long-term goal is to decipher how the recently identified BrS risk alleles are functionally associated with this cardiac disease. Here, by using the fly model, we aimed at determining which of the 56 genes near the identified loci are involved in cardiac physiology using high-throughput functional screening. IRX 5, IRX3, TBX20 and GATA4 were not included in this study, as their roles as cardiac transcription factors have already been described^17,18^. According to the DRSC Integrative Ortholog Prediction Tool (DIOPT: https://www.flyrnai.org/diopt), among the human genes identified as being potentially linked to BrS-risk alleles, 38 (70%) are orthologs to 44 fly genes (DIOPT rank High or Moderate, score >2). (Figure 1A, Table S1). Cardiac-specific gene knockdown (KD) was performed using publicly available transgenic flies expressing double-stranded RNA (dsRNA) constructs under Upstream Activating Sequences (UAS) control, that specifically interfere with gene expression. Transgenic lines were from both the VDRC (Vienna Drosophila Resource Center https://shop.vbc.ac.at/vdrc_store/) and the DRSC-TriP (https://fgr.hms.harvard.edu/trip-rnai-fly-stocks) collections. Among the 44 fly genes selected, we found that 33 genes had 2 independent available KD lines, 10 genes had only one KD line available and 1 gene did not have specific dsRNA line available in the public collections. Overall, 70 dsRNA transgenic lines, targeting 43 genes, were used in this study (Table S2). We used the heart specific Hand>Gal4 driver^19^ to inhibit gene function in the whole cardiac system, including cardiomyocytes and associated non-contracting pericardial cells (Figure 1B). After adult emergence, males and females were aged separately to one week and used for *in vivo* imaging of heart function on intact flies as previously described^10^.

**Figure 1:**
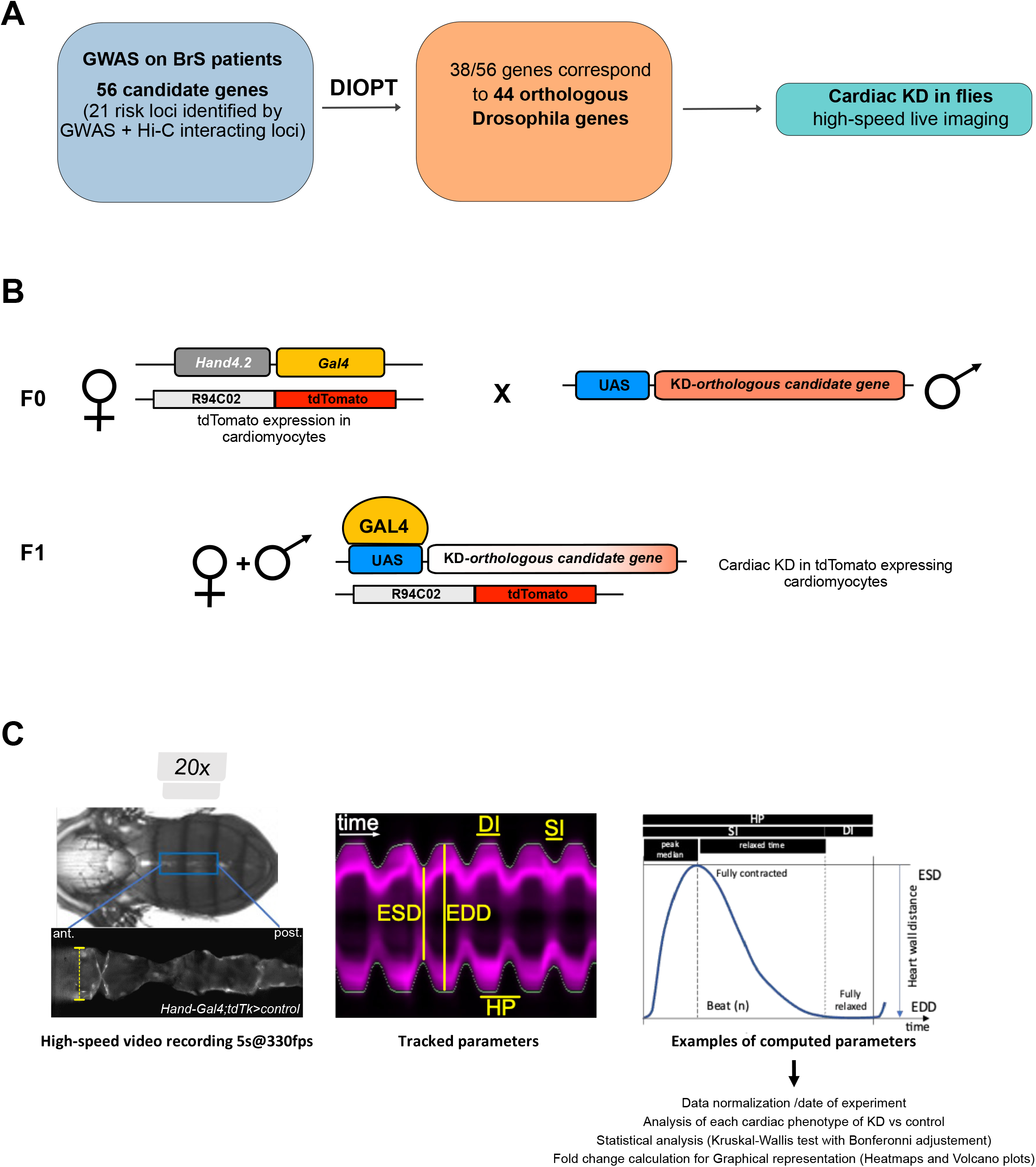
High-throughput cardiac functional validation of Drosophila orthologues associated to human BrS. A) Selection scheme of fly orthologues with DIOPT, to be investigated by cardiac KD. B) Crosses realized to invalidate cardiac gene expression. C) Left: intact anesthetized 1-week Hand>Gal4/+ adult female fly expressing tdTomato in the heart. Middle: M-mode record generated at the position indicated by dotted line, showing end diastolic (EDD) and end systolic (ESD) diameters, diastolic interval (DI), systolic interval (SI) and heart period (HP). Right: representation of one heartbeat, and associated computed parameters.

Automatic processing of video captions allowed to monitor cardiac parameters of individual flies. As illustrated on kymograms (M modes, Figure 1C), to characterize heart rhythm, we focused on heart period (duration of one heartbeat, HP) and on the duration of systolic interval (SI) and of diastolic interval (DI) separately. Several parameters were also evaluated as a measure of rhythm variability: the median average deviations of HP, SI and DI (MAD-HP, MAD-SI and MAD-DI) as well as Arrhythmia Index (AI), corresponding to the standard deviation of HP normalized by HP mean. Kymograms are also used to measure heart diameters at full contraction (End Systolic Diameter, ESD) and full relaxation (End Diastolic Diameter, EDD). Both ESD and EDD are then used to calculate the fractional shortening (FS = EDD-ESD/EDD) with estimates the contraction efficiency of the heart. Each UAS>dsRNA collection being established in a specific genetic background, all experiments were performed together with control crosses using corresponding genetic background, allowing for normalization for both genetic background and day of experiment (see M&M).

The effect of gene knockdown on cardiac parameters were compared to controls and corresponding fold changes were calculated. Kruskal-Wallis test was used to estimate the statistical significance of the fold changes and Bonferroni correction allowed adjustment for multi-testing. Fold changes values and significance were visualized on volcano plots for detailed description of the results per sex and per phenotype (Figure 2 and Figure S1) and on heatmaps for global assessment of knockdown effects on phenotypes (Figure 3).

**Figure 2:**
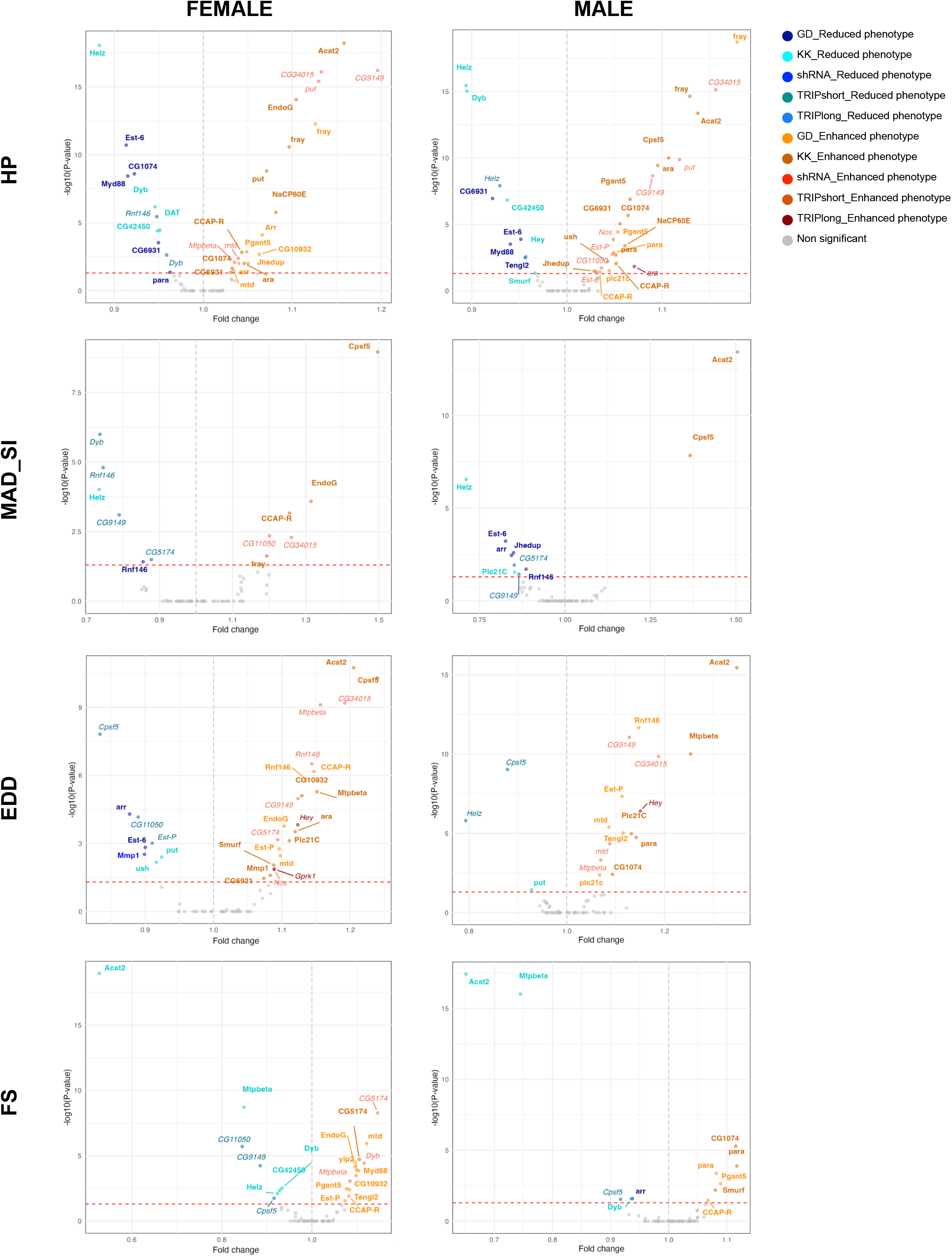
Volcano plots showing quantified phenotypes resulting from all performed gene knock down in the heart. Results obtained in females and males are shown for heart period (HP), median average deviation for SI (MAD_SI), end diastolic diameter (EDD) and fractional shortening (FS). Fold changes and adjusted P-values (<0.05) are indicated. Colors correspond to the different collections of transgenic lines. Shades of blue show reduced phenotypes compared to respective controls, and shades of red/orange represent enhanced ones.

**Figure 3:**
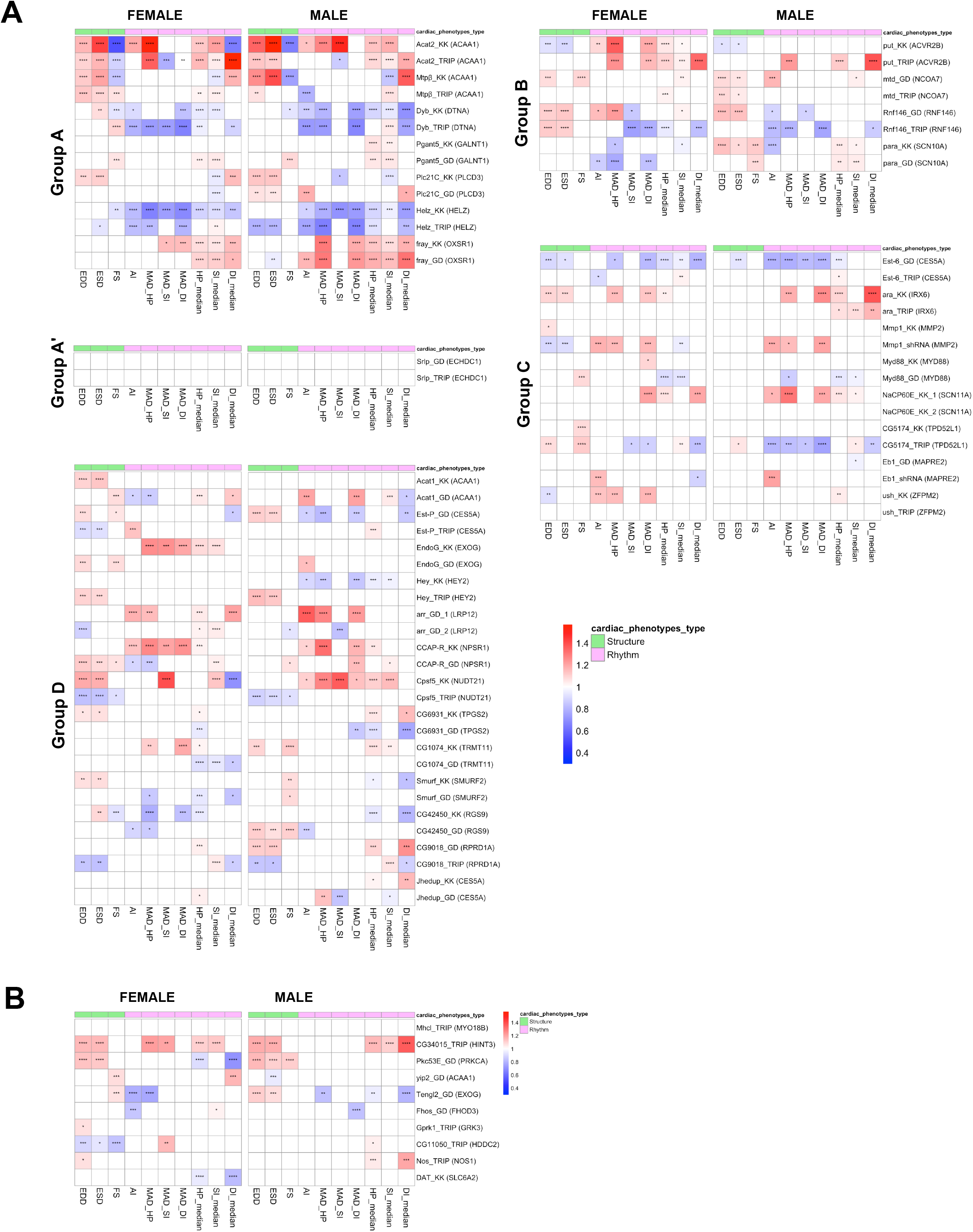
Unclustered Heat maps showing quantified phenotypes resulting from all cardiac specific gene knock down performed. Cardiac phenotypes for structure (green) and rhythm (pink) are represented. For each group, female and male phenotypes are shown. Fold changes (red to blue ladder) and adjusted P-values are indicated (* p<0.05 ; ** p<0.01 ; *** p<0.005 ; **** p<0.0001).

Overall, all dsRNA lines led to at least one significant phenotypic difference with respect to its control in either females or males (Figure 3). This illustrates the high sensitivity of cardiac phenotypes detection that was used. HP was the most affected phenotype, being modified in almost half of the lines analyzed (47% and 52% of lines were associated with HP modification in females and males respectively, see Table 1 and Figure 2), while MAD-SI was the least sensitive phenotype (17% of lines exhibited modified MAD-SI in females, 14% in males, see Table 1 and Figure 2). Though most cardiac parameters were equally modified in both sexes, EDD and FS exhibited sex-biased changes as a result of gene KD (44% of KD lines displayed modified EDD in females, compared to 26% in males, 31% of KD lines displayed modified FS compared to 16 % in males, see Table 1 and Figure 2).

**Table 1:** Proportion of lines with significant differences of indicated phenotypes, by sex.

|  | EDD | ESD | FS | AI | MAD-SI | MAD-DI | MAD-HP | HP | SI | DI |
| --- | --- | --- | --- | --- | --- | --- | --- | --- | --- | --- |
| Females | 0.44 | 0.35 | 0.31 | 0.29 | 0.17 | 0.31 | 0.34 | 0.47 | 0.43 | 0.32 |
| Males | 0.26 | 0.31 | 0.16 | 0.35 | 0.14 | 0.31 | 0.32 | 0.52 | 0.41 | 0.35 |

Given the sensitivity of the assay, to confidently identify genes whose function is cell autonomously required for heart functioning, we first focused on the ones that were knocked down by two independent dsRNA lines (Figure 3A). Genes were grouped into four classes (A-D, Figure 3A) depending on the consistency of the cardiac defects observed in both sexes and with both dsRNA lines.

Group A is composed of 7 genes that induced similar phenotypes in both females and males when knocked down by each of the two lines. These are Acat2 (corresponding to human *ACAA1* gene), *Mtpβ* (*ACAA1*), *Dyb* (*DTNA*), *pGANT5* (*GALNT1*), *plc21C* (*PLCD3*), *Helz* (*HELZ*) and fray (*OXSR1*). Congruent results obtained in both sexes and the similar phenotypic characteristics resulting from the two lines are strong arguments for considering those genes as being involved in heart functioning. Corresponding human genes may therefore be given high priority for further investigations.

*Group A’* is composed of one gene (*srlp* (*ECHDC1*)) that led to lethality in both males and females when knocked down in the heart with the two dsRNAi lines used. This presumably reveals that this gene fulfills an essential - but yet unknown - function in the fly heart.

*Group B* is composed of 4 genes whose knockdown gave similar phenotypes with both dsRNA lines in one sex but weak or different phenotypes in the other sex. This was the case for put (*ACVR2B*) with similar phenotypes for both KD lines in females (increased heart period and arrhythmia) but inconsistent phenotypes in males, for *mtd* (*NCOA7*) whose knockdown was associated with dilated heart (increased EDD and ESD) in males but not in females, for *Rnf146* (*RNF146*) with dilated heart and reduced SI variability in females but reduced arrhythmia in males and lastly for para (*SCN5A/SNC10A*) whose dsRNA lines lead to increased heart period (increased HP and SI) and increased fractional shortening (FS) in males but reduced heart period variability (reduced MAD-HP) in females. This may point to sexual dimorphism in gene’s activity with respect to heart functioning, or, alternatively may be due to weak dsRNA efficiency in one of the two sexes. All 4 genes may deserve further attention in mammalian models, mostly regarding the stronger incidence of BrS observed in men.

Group C is composed of 8 genes (*ara* (*IRX6*), *Est-6* (*CES5A*), *Mmp1* (*MMP2*), *NaCP60E* (*SCN11A*), *Myd88* (*MYD88*), *Eb1* (*MAPRE2*), *ush* (*ZFPM2*) and CG5174 (*TPD52L1*)) that led to consistent phenotypes among sexes when knocked down with one dsRNA but led to weaker phenotypes when using the second dsRNA line. Those genes may also deserve further investigation: weak effects observed with one dsRNA line is most probably due to the relative inefficiency of one dsRNA line in our cardiac test. Importantly, knocking down *ush* (*ZFPM2*), with one of the 2 available lines resulted in major morphological and functional defects. In addition, while the heart in *Drosophila* is in close contact with the dorsal cuticle - enabling *in vivo* imaging thanks to the tdTK reporter used here-this knock down led to hearts detached from the cuticle and floating in the abdominal cavity, preventing video capture. The second *ush* dsRNA line used may be much weaker and led to increased arrhythmia specifically in females. The observed phenotypes therefore prompt further investigations of *ZFPM2* function in mammalian models.

Group D is composed of 13 genes (*Est-P (CES5A), Acat1(ACAA1), EndoG (EXOG), Hey (HEY2), arr (LRP12), CCAPR (NPSR1), Cpsf5 (NUDT21), CG6931 (TPGS2), CG1074 (TRMT11), Smurf (SMURF2), CG42450 (RGS9), CG9018 (RPRD1A), Jhedup (CES5A)* that led to cardiac phenotypes using both dsRNA lines but with different or sometimes opposite phenotypes from one line to the other. The discrepancy observed among RNA lines targeting the same genes prevent firm conclusion to be drawn regarding this group. For instance, Hey (HEY2) knock down lead to cardiac hypertrophy in both males and females when driven by one dsRNA transgene, but to male specific reduced arrhythmia, HP and SI when driven by the other one. Given the known involvement of HEY2 in BrS syndrome (ref), we expected reproducible cardiac functional defects following the knock down of its Drosophila orthologue. This discrepancy may be due to off target effects of one of the two lines and therefore point out a limitation of our approach.

Among the 10 remaining candidate genes for which one dsRNA line was tested (Figure 3B), one (*mhcl* (*MYO18B*)) lead to lethality when driven by Hand>Gal4 and two (*CG34015* (*HINT3*) and *pkc53E* (*PRKCA*)) lead to similar phenotypes among females and males. All 3 genes should therefore be considered for further investigations. The other 7 genes were characterized by weak and/or inconsistent phenotypes among sexes and, in the absence of alternative RNAi lines to test the robustness of those phenotypes, may not be given high priority.

Based on their available genetic and molecular function, we discuss below the possible involvement in cardiac functioning of genes identified in groups A and B.

- *Acat2 (Acetyl-CoA acetyltransferase 2)* and *Mtpβ (Mitochondrial trifunctional protein β subunit)* encoding acetyl-CoA C-acetyltransferase are both involved in fatty acid (FA) beta-oxidation. They are predicted to be localized in peroxisome and the mitochondrion, respectively. When knocked down in the heart both genes led to cardiac hypertrophy (increased EDD and ESD), which was associated to reduced fractional shortening (FS). In addition, *Acat2* knockdown was associated with a reduced heart rate (increased HP and SI) and increased arrhythmia. Fatty acids being the main source of energy supply in adult contractile cardiac cells, reduced beta oxidation directly affects acetyl-CoA levels and might therefore directly impact heart functioning by impinging on ATP production in the cardiomyocytes. A prolonged reduction of this energy pathway might have side effects on glucose homeostasis itself, by altering the catabolic equilibrium dependent on mitochondrial function that may lead to cardiac dysfunctions.
- *Dyb (Dystrobrevin)* encodes a structural constituent of muscle cells, which is part of the Dystrophin-associated glycoprotein complex. The Dystrophin glycoprotein complex is a plasma transmembrane complex that links the actin cytoskeleton to the extracellular matrix and is important for maintaining the integrity of skeletal and cardiac muscle cells. In flies, *Dystrophin (Dys)* loss of function alleles were associated to increased arrhythmia and dilated cardiomyopathies^20^. Our data established that *Dyb* heart specific knockdown was associated to reduced arrhythmia and increased heart rate (decreased heart period) but did not consistently lead to structural defects. The observed discrepancy between *Dyb* and *Dys* phenotypes could indicate that *Dyb* could fulfill specialized and independent function(s) within the Dystrophin complex. It alternatively may reveal different effects resulting from the nature of the genetic alteration analyzed in each study (genomic deficiency leading to organismal loss of function for *Dys* versus heart-specific RNAi mediated knockdown for *Dyb*). Of note, missense mutations of *Dyb* mammalian orthologues (DTNA) were associated to congenital forms of left-ventricular non compaction cardiomyopathies^21^.
- *pGant5* (*Polypeptide N-Acetylgalactosaminyltransferase 5*) encodes a member of the family of glycosyltransferases (GalNAc-Ts). It initiates mucin-type O-glycosylation by catalyzing the transfer of a GalNAc sugar to the hydroxyl groups of serine and threonine residues in secreted or membrane-bound proteins, thereby influencing extracellular matrix (ECM) composition and signaling pathways. In mice, GALNT1 is associated to abnormal valve development and subsequent cardiac dysfunctions^22^. GALNT1-mediated glycosylation also controls the balance between active and inactive brain natriuretic peptide (BNP), triggering increased proportion of inactive proBNP secreted by the heart when dysregulated, and impairing the compensatory actions of BNP during the progression of heart failure^23^. In flies, our results showed that pGant5 knockdown was specifically associated to increased heart period (and specifically to increased systolic interval), probably mediated by ECM and/or signaling pathways modifications.
- *plc21C (Phospholipase C at 21C)* encodes a phospholipase C beta that is involved in the regulation of several signaling pathways, notably in Wnt/TCF signaling pathway. In mice PLCD1 and D3 appeared to have redundant function(s) and their loss of function was associated to cardiac fibrosis, dilation, and hypertrophy^24^ We showed that *plc21C* knockdown in flies is also associated to dilated hearts (in both systole and diastole). It additionally reduced systolic interval, with no incidence on heart rate, probably due to a compensatory increase in diastolic interval.
- *Helz (Helicase with zinc finger)*. Fly and human HELZ proteins were found associated with mRNA decay, throughout their interaction with the CCR4-NOT deadenylase complex^25^. Interestingly, silencing of CCR4-NOT components in adult Drosophila was previously shown to result in dilated cardiomyopathy, while heterozygous *NOT3* knockout mice showed impairment of cardiac contractility and increased susceptibility to heart failure^26^. In humans, a common variant in the *NOT3* gene has been associated to altered cardiac QT intervals^26^. Our results demonstrated that fly cardiac specific knockdown of *Helz* was associated to accelerated heart rate (decreased HP, SI and DI) and reduced arrhythmia. These phenotypes suggest that the HELZ-mediated mRNA decay process may be involved in heart rhythm regulation.
- *fray (frayed)* encodes a serine threonine protein kinase whose activation is mediated by the Wnk kinase. It regulates the activity of various ion transporters and channels. *fray* is involved in a large panel of biological processes, from K^+^ buffering linked to regulation of seizure susceptibility^27^ to mitochondrial energy metabolism^28^ and regulation of Wnt signaling pathway^29^. In humans, OXSR1 has been associated to early cardiac development^30^ but, to date, no study established its involvement in adult heart functioning. Interestingly heart specific knockdown of fray in flies resulted in reduced heart rate (increased HP, SI and DI) and increased arrhythmia.
- *srlp* human orthologue ECHDC1 (ethylmalonyl-CoA decarboxylase) triggers decarboxylation of ethyl- or methyl-malonyl-CoA. It is involved in branched-chain fatty acid (FA) metabolism, by preventing the formation of potentially deleterious ethyl-branched FAs^31^. There is no known function for ECHDC1 in the heart. Importantly, our results showed that heart-specific srlp knockdown using both RNAi lines, leads to lethality in flies, strongly suggesting that ethyl branched FAs may have deleterious effects on cardiac function.
- *put (punt)* encodes the transforming growth factor beta (TGFβ) type II receptor that acts in both Dpp/BMP and Activin signaling pathways. Our previous studies have identified both signaling pathways as synergistic interactors for heart rhythm and cardiac diameters regulations. Indeed, loss of function mutations of components of these pathways were associated with reduced cardiac diameters and increased HP, SI and DI^10^. Of note put heart specific knockdown was associated with reduced ESD and EDD in both males and females, but with increased heart period and arrhythmia specifically in females.
- *Rnf146 (Ring finger protein 146)* encodes a ubiquitin protein ligase involved in proteasome ubiquitin mediated protein degradation. As such, it is involved in Wnt pathway regulation both in flies^32^ and in mammals. Regarding its function in the heart, in mammals, it alleviates myocardial ischemia/reperfusion injury by promoting Dapk1^33^ degradation and protects against oxidative stress-mediated cardiac injury by targeting the degradation of PARP-1 ^34^. In flies, cardiac specific knockdown was associated to dilated heart and reduced SI variability in females but increased arrhythmia in males.
- *mtd (mustard)* encodes a nuclear receptor co-activator regulated by the fly steroid hormone Ecdysone. Its orthologue NCOA7 has been demonstrated to be involved in resistance to oxidative stress in several biological processes^35,36^. mtd knockdown was associated with cardiac dilation specifically in males, pointing to potential sex specific involvement of the gene.
- *para (paralytic)* encodes the α-subunit of voltage-gated sodium channels (SCN5A/SCN10A). While no function for Na+ channels in cardiac functioning was reported to date in flies, our results demonstrate that para knockdown in the heart lead to reduced heart rate (increased HP and SI) specifically in males, thus pointing to a putative sex-biased control of heart rhythm.

## Conclusions

The highly complex genetic architecture of the Brugada syndrome calls for an in-depth analysis of candidate genes associated to the pathology. Here, by implementing a high throughput screening of orthologues of human genes in the fly model, we were able, on a large scale, to analyze the effects of heart specific invalidation of genes on cardiac function *in vivo*. Our results will allow prioritization of genes for in depth analysis in mammalian models, such as human iPSC derived cardiac organoids. Identifying the functional consequences of the identified risk alleles might allow to identify new markers of arrhythmic risk and possibly also new therapeutic targets.

## ACKNOWLEDGMENTS

We thank the WIRES consortium for helpful discussions and comments on the manuscript. We thank Frederic Gallardo and Tahagan Titus for their technical assistance in fly food cooking. We also thank the Bloomington Stock Center and the Vienna Drosophila RNAi Center for fly stocks. Funding is from ANR (WIRES, grant n°ANR-22-CE17-0051-03 to N.A. and L.P.), and Amidex (grant n°AMX-21-PEP-020 to N.A.), INSERM and Aix-Marseille University (AMU).

## CONTRIBUTIONS

Conceptualization: L.P., N.A., J.B., F.C., Methodology: S.K., L.P., N.A., Experimental investigation: S.K., Data analysis: S.K., L.P., N.A., Visualization: S.K., N.A., Supervision: L.P., N.A., Writing-Original Draft: N.A., L.P., corrections of the original Draft: F.C., N.G., Secure Funding: N.A., L.P.

## MATERIALS AND METHODS

### 1. Selection of fly orthologues of human genes associated to BrS syndrome

Drosophila orthologues of the 56 human genes located near loci associated with Brugada Syndrome (BrS), were identified and selected using the DIOPT tool (DRSC Integrative Ortholog Prediction Tool, (https://www.flyrnai.org/diopt). Only orthologues with a DIOPT score above 2 - indicative of moderate to high confidence - were retained. 44 Drosophila genes corresponding to 38 human genes were finally selected for cardiac functional screening.

### 2. Selection of dsRNAi lines to be tested for functional screening and crosses for heart specific knock down

Transgenic lines selection was guided by the UP-TORR database (https://www.flyrnai.org/up-torr/), prioritizing constructs with high knockdown efficiency and minimal off-target effects. When possible, two independent dsRNAi lines per gene were used to ensure reproducibility. Lines targeting the selected Drosophila genes orthologues were obtained from the Vienna Drosophila Resource Center (VDRC) and the DRSC collection (TriP) resourses. Cardiac specific knockdown was achieved using the Hand4.2-Gal4 driver crossed to UAS-RNAi lines, along with tdTomato (R94C02::tdTomato)^37^ reporter expressed in cardiomyocytes. F1 progeny expressed both RNAi and tdTomato in the heart, allowing functional knockdown and cardiac live imaging.

Given the strong male predominance of Brugada Syndrome in humans, both male and female flies were analysed. To reflect the adult-onset nature of BrS, cardiac function was assessed in 7 days-old adult flies.

### 3. Fly Husbandry and Stocks

All Drosophila stocks were reared at 18°C on standard food, while experimental crosses were performed at 25°C with a 12/12hours light/dark cycle and 50% humidity. Standard food contained per liter: 10g agar, 90 g corn flour, 90 g inactivated yeast extract, and 3.75 g Moldex (in 70% Ethanol). F1 progeny flies were maintained at 25°C on 2/12hours light/dark cycle for 7 days before analysis.

### 4. Fly heart live recording and analysis

Live cardiac imaging was performed on intact adult flies anesthetized with FlyNap (Sordalab) as previously described^10^. Following preliminary tests to optimize both dosage and exposure time to Flynap, in order to avoid significant effects on heart rhythm or contractility, a 1min anesthesia was chosen and flies allowed to recover for 5 minutes, a period sufficient to restore stable cardiac activity. Flies immobilized on coverslips using a UV-curing glue (Norland Optical Adhesive, #NOA61) are imaged under a 20× dry objective (Zeiss Axioplan microscope). Cardiac activity in the A2–A3 abdominal region of heart tubes expressing tdTomato reporter, was recorded for 5 s (330 frame per second), using a high-speed CMOS camera (Orca Flash4.0, Hamamatsu Photonics) and HCI imaging software.

For each recording, M-mode kymographs were generated, and key cardiac parameters were calculated using a custom R pipeline (https://github.com/gvogler/FlyHearts-tdtK-Rscripts) developed by G Vogler. Extracted parameters included heart period (HP), systolic interval (SI), diastolic interval (DI), end diastolic diameter (EDD), and end systolic diameter (ESD). Derived metrics, including heart rate (1/HP) and fractional shortening [(EDD − ESD) / EDD], were also computed (Figure 1C). Approximately 40 flies per sex and per dsRNAi line were analyzed.

To control for technical variability, all imaging was performed within standardized time windows post-recovery and at fixed times of day. Males and females were tested on the same day, along with background-matched controls. Day and sex specific normalization was performed by scaling raw values to the mean of the control for each group. Normalized values were used for all subsequent statistical analysis.

### 5. Data Visualization

Two complementary visualization strategies were implemented in R to assess cardiac phenotypes by gene and sex. Clustered heatmaps were generated using the “pheatmap” package, displaying fold changes across 10 cardiac parameters in males and females. Parameters were grouped by functional class (structural vs rhythmic), and adjusted p-values were overlaid on heatmap cells for clear overview of effect intensity and significance. Volcano plots were generated using “ggplot2” package, plotting log_2_ fold changes against −log_10_(adjusted p-values) for each phenotype and sex. RNAi lines were color coded by source (VDRC or TriP) and direction of effect (increase or decrease). These visualizations supported the classification and prioritization of candidate genes for further functional validation in mammalian models.

### 6. Statistical Analysis

All statistical analyses were conducted in R. Raw data were normalized per day and sex relative to the mean of matched control groups. Outliers were removed using the interquartile range (IQR) method for each group. For each phenotype, fold changes were calculated as the ratio between RNAi and control means. Group differences were tested using Kruskal-Wallis non-parametric tests, followed by Bonferroni correction for multi-testing. Adjusted p-values were categorized as follows: ****⍰< ⍰ 0.0001, ***⍰< ⍰0.001, **⍰< ⍰0.01, *⍰< ⍰0.05, NS⍰≥⍰0.05.

All results were compiled into summary tables reporting fold changes, adjusted p-values, direction of effect (increase, decrease, no change), and were visualized using heatmaps and volcano plots as described above.

We used FlyBase (release FB2025_02) to update information on candidate genes, their function and described phenotypes.

## Supplemental Figures and Tables

**Figure S1 related to Figure 2: Volcano plots** showing quantified phenotypes resulting from all performed gene knock down. Results obtained in females and males are shown for end systolic diameter (ESD), diastolic interval (DI), systolic interval (SI), arrhythmia index (AI), median average deviation for HP (MAD_HP), and for diastolic interval (MAD_DI). Fold changes and adjusted P-values<0.05 are indicated. Colors correspond to the different collections of transgenic lines. Shades of blue show reduced phenotypes and shades of red/orange enhanced ones.

**Table S1:** List of human genes identified in GWAS analysis/ DIOPT score for all potential fly orthologues with indication of the ones tested in this study.

**Table S2:** List of the independent transgenic dsRNAi lines tested, corresponding to every fly orthologue identified. Lines belong to the VDRC and BDSC public collections.

## Data sets

Raw data of cardiac imaging

Normalized data presented in the results

## R-scripts for data analysis and graphical representations

All scripts are available on GitHub [https://github.com/sallouha/BrS_paper].

