## Supplementary figures and images for "Exploring cardiac phenotypes of Drosophila orthologues of human genes associated to Brugada Syndrome (BrS)"

### Supplemental Figure 1

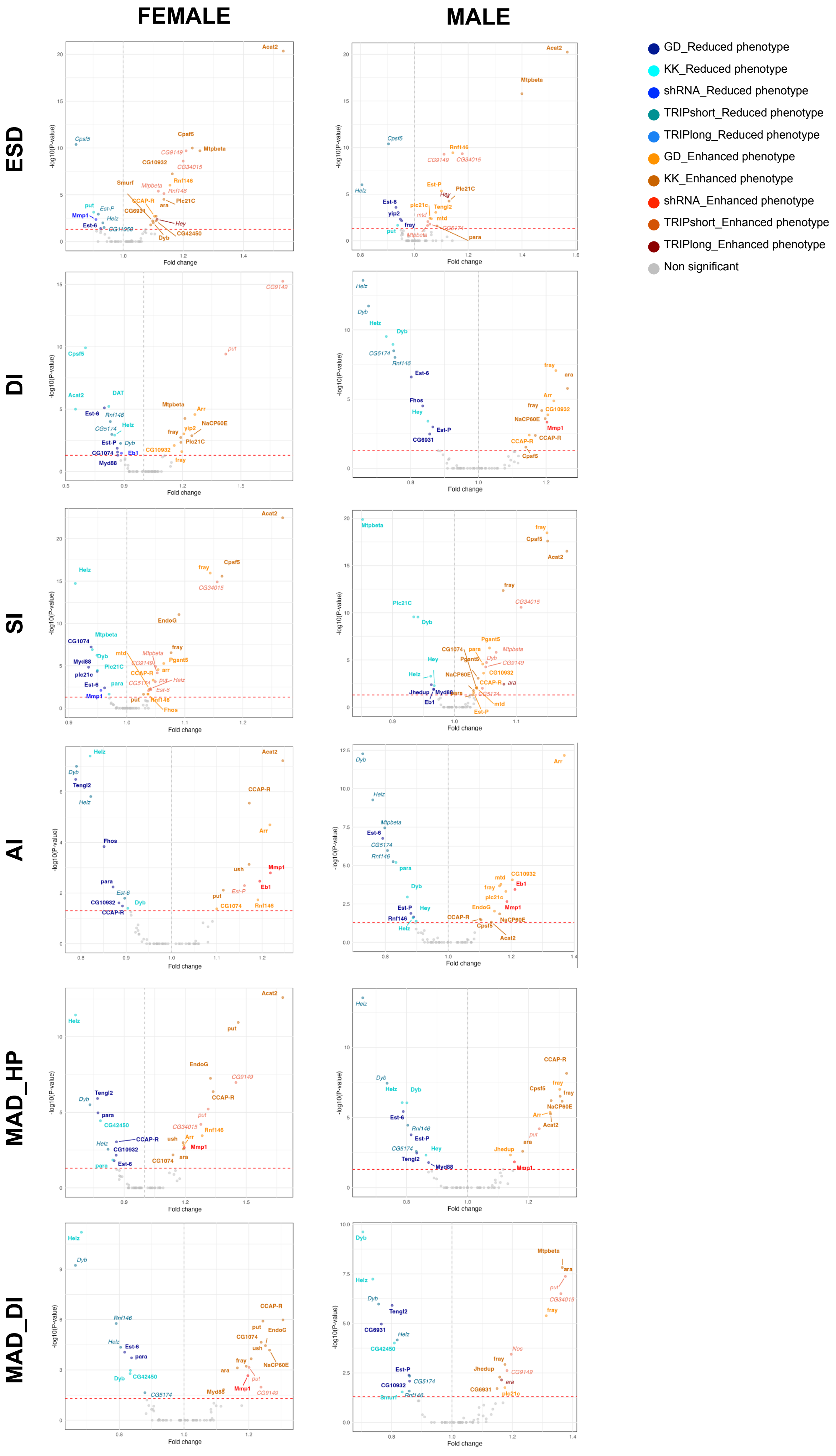
